# Directionality bias in T/A cloning

**DOI:** 10.64898/2026.02.11.705383

**Authors:** Valeria Dountcheva, Athanasios Bubulya, Labib Rouhana

## Abstract

T/A cloning is a popular method for generating recombinant DNA plasmids. This method relies on single A:T nucleotide base pairs between PCR product ends and vector. Theoretically, the directionality of insert ligation with relation to the vector is random. However, we have continuously observed directionality bias using the pGEM-T Vector System for T/A cloning in a Course-based Undergraduate Research Experience (CURE). Cloning of over 400 inserts has shown directional bias higher than 74% (*p-*value < 0.0005) “sense” to the T7 promoter of the vector. Awareness of biased insertion in our applications reduces time and cost in cloning and downstream analyses.

## Description

*Taq* DNA Polymerase deposits a non-templated adenosine at the 3’end of DNA strands (Clark, 1988; Williams, 1989). This 3’-end adenosine “overhang” is exploited for rapid and effective insertion of PCR products into plasmid vectors that carry single thymidine overhangs during “T/A cloning”(Holton and Graham, 1991; Kovalic et al., 1991; Marchuk et al., 1991; Mead et al., 1991). T/A cloning is favorable to blunt-end ligation due to the increased rate of success (Holton and Graham, 1991; Liu et al., 2018). In theory, the directionality of ligation between the insert and vector in T/A cloning is random. However, directional insertion is often desirable during cloning to facilitate downstream applications such as expression of proteins and generation of riboprobes in high throughput analyses. While directional cloning is a time-consuming technique reliant on thoughtful design and processing with multiple restriction enzymes, we have observed a strong directional bias using the pGEM-T Vector System for T/A cloning. In this report, we present evidence obtained from several years of cloning using the pGEM-T vector system in our Course-based Undergraduate Research Experience (CURE) that demonstrates statistically significant bias in the directionality of inserts.

A bias in directionality of partial cDNA sequence inserts in T/A cloning events was observed repeatedly over 6 years of activities in a CURE using the pGEM-T vector (Figure 1A). In each CURE project, partial cDNA amplicons corresponding to approximately 400-600 bps of planarian (*Schmidtea mediterranea*) sequence were inserted into pGEM-T and individual clones validated through Sanger sequencing. Quantification of insert orientation with respect to the T7 and SP6 promoters flanking the insertion site, revealed that 75% of the insertions generated for Project 1 were positioned in the sense orientation relative to the T7 promoter (Figure 1B).

**Figure 1.**
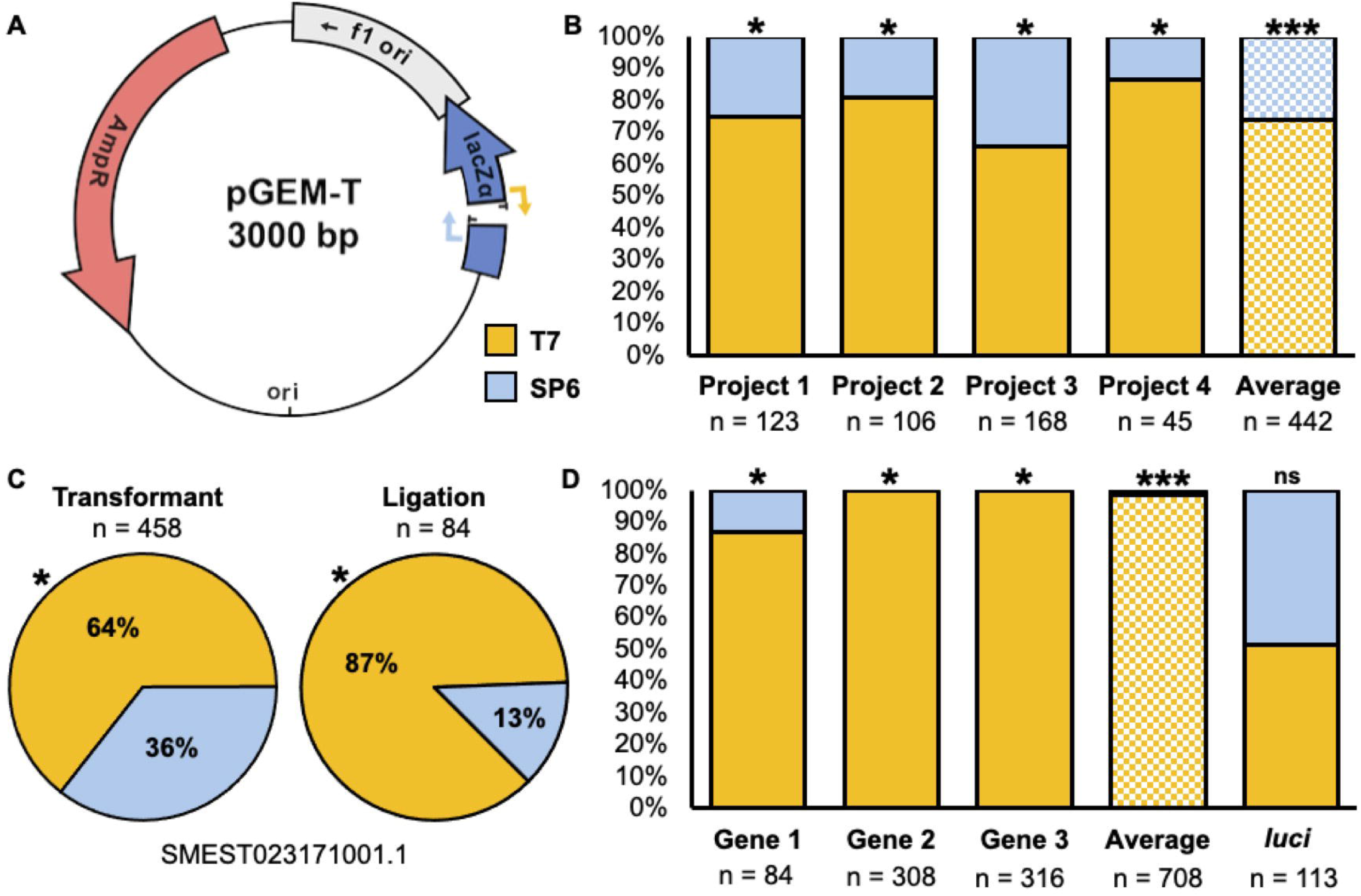
Orientation bias in insertion of DNA fragments during T/A cloning using the pGEM-T vector. **A)** Plasmid map of pGEM-T illustrates the position of T-overhangs used in T/A cloning and flanking T7 and SP6 promoters. **B)** Percentage of cDNA insertion events in the sense orientation relative to the T7 promoter (yellow) and the SP6 promoter (blue) in constructs generated over the course of four CURE projects. The average across the projects is shown (checkered graph). **C)** Measured distribution of insert orientation from a cloning attempt of SMEST023171001.1 (*pumilio* homolog) cDNA fragment assessed after the ligation and transformant selection steps. **D)** Bar graphs display orientation distribution of inserts after the ligation step from the cDNA fragment in (C; Gene 1) and two additional genes (SMEST004045004.1, Gene 2; and SMEST012773001.1, Gene 3), as well as their average (checkered bar). Orientation of Control Insert DNA (*luci*) included in the pGEM-T Vector System after the ligation step is also shown. Asterisks show statistical significance according to (* = p < 0.005) the chi-squared test and (*** = *p* < 0.0005) unpaired Student’s *t-*test.

Enrichment of sense orientation relative to the T7 promoter was also observed in subsequent projects as follows: 81% for the Project 2, 65% for Project 3, and 87% for Project 4 with each found to be statistically deviant from random insertion using the chi-squared test. On average, 74% of the inserts were positioned sense to the T7 promoter in the 442 constructs generated by over 100 separate individuals (Figure 1B). Statistical analysis using Student’s *t-*test revealed that the overall deviation in directionality of insertion is significantly deviant from the 50% distribution expected for T/A cloning (*p* < 0.0005).

To understand how this bias might be generated, we looked at the structure of the pGEM-T vector. The insertion site for T/A cloning lies within the *LacZ*α reporter gene used in blue/white colony screening in the pGEM-T system. *LacZ*α is transcribed from a promoter upstream and in the same orientation as the SP6 promoter, and opposite orientation from the T7 promoter, flanking the insertion site. White colonies are picked in cloning attempts because these represent successful insertion of amplicons within *LacZ*α and therefore disruption of a functional *LacZ*α product. We considered the possibility that insertions of small portions of ORF in the sense orientation relative to *LacZ*α could result in a functional enzyme and therefore generate blue colonies representing false negative events. Therefore, we hypothesized that the bias observed in insert orientation may be an artifact of blue/white screening. To tease this apart, a fragment corresponding to a planarian *Pumilio* homolog (SMEST023171001.1) was amplified from cDNA and ligated into pGEM-T. A portion of the ligation reaction was transformed into *Escherichia coli* JM109 cells, while another portion of the ligation reaction was used directly as template to sequence across the insertion site between the T7 and SP6 promoter using third generation long read sequencing (PCR-EZ, Azenta). 64% of insertion events sequenced from white colonies were sense orientation relative to the T7 promoter in this experiment (458 reads, Figure 1C). Unexpectedly, sequences obtained directly from the ligation reaction also showed a strong bias in sense orientation relative to the T7 promoter (87% of 84 reads; Figure 1C).

Ligation reactions were sequenced for two more planarian cDNA amplicons (SMEST004045004.1 and SMEST012773001.1), both of which revealed 100% insertion events in the sense orientation relative to the T7 promoter (624 reads, Figure 1C). Sequencing results from each of the three ligation reactions deviate significantly from the expected random directionality of insertion according to chi-squared test. Together the results from sequencing insertion events prior to transformation into *E. coli* showed a 98% average of insertion of planarian ORF gene fragments in the sense orientation relative to the T7 promoter (*p* < 0.0005, Figure 1D). Finally, to investigate whether insert directionality bias is observed when cloning DNA fragments obtained from other sources, we sequenced a ligation reaction using the control insert provided in the pGEM-T Vector System, which is a 542 bp fragment of the firefly *luciferase* gene. Ligations using the control insert showed no orientation bias, as would be expected from random insertion events (Figure 1D). These results indicate that something about the cDNA fragments selected, or the mechanism by which they are derived, is driving the bias in directionality observed during T/A cloning of planarian cDNA fragments.

Further investigation is needed to determine the mechanism driving the orientation bias observed at the ligation step during T/A cloning using the pGEM-T Vector System. Perhaps coding sequences in our model, the planarian *S. mediterranea*, are composed of nucleotide combinations that cause preferential structural compatibility with one end of the pGEM-T vector cloning site. For a few inserts (*e*.*g*., *Smed-nanos* homolog, SMEST018169001.1), we saw strong orientation bias sense to the SP6 promoter, which is the opposite direction from the vast majority of our clones. Another possibility is that the tool used for designing primers in our cloning efforts (Primer3; Koressar and Remm, 2007; Untergass et al., 2012) integrates an unapparent difference in composition of the forward primer from the reverse primer (in our case, the forward primer is in the sense orientation). Comparing 96 primer sets revealed that the forward primers had 10% more purines than their reverse counterparts, which is a statistically significant difference (*p* = 1.94 × 10^-06^). Other parameters of primer composition, such as GC content, annealing strength, annealing temperature, and length were all comparable. Regardless of the mechanism, awareness of this bias has allowed us to streamline cloning of fragments utilized for generation of riboprobes in our mid-scale screens. It will be interesting to see if others observe the same phenomenon using sequences from other organisms or in different T/A cloning systems.

## Methods

### cDNA amplification and cloning

RNA was extracted from *Schmidtea mediterranea* sexually mature hermaphrodites using TRIzol™ Reagent (Thermo Fisher Scientific, Waltham, Massachusetts). cDNA was generated using the GoTaq 2-Step RT-qPCR System (Promega, Madison, Wisconsin) with 1 □g of total RNA as template and a mix of random and oligo(dT) primers following the manufacturer’s instructions. Gene-specific primers targeting 400 –600 bps of cDNA coding regions (whenever reference sequence made this length range possible) were designed using the Primer3 web tool (https://primer3.ut.ee/; (Koressaar and Remm, 2007; Untergasser et al., 2012). The following parameters were specified during primer design: T_m_: 61-63 °C (optimal 62 °C); GC content: 35-65% (optimal 50%); primer size: 18-26 nts (optimal 22 nts); GC clamp: 1 nt. The remaining parameters were used at the default settings. cDNA amplicons were generated using GoTaq® Long PCR Master Mix (Promega, Madison, Wisconsin) with an initial denaturation at 95 °C for 2 minutes, followed by 40 cycles of denaturation at 95 °C for 30 seconds, annealing at 56 °C for 30 seconds, and extension at 72 °C for 1 minute, and a final extension at 72 °C for 5 min.

Amplicons were then cleaned up using the DNA Clean & Concentrator-5 kit as per manufacturer instructions (Zymo Research, Tustin, California) and eluted in 20 □L of nuclease-free water. A volume of 1 □L of clean amplicon was ligated into the pGEM-T vector using the Standard Reaction volume as per manufacturer’s instructions (Promega, Madison, Wisconsin) and 3 □L of ligation reaction were transformed into 30 □L JM109 cells. Plasmids were purified from liquid Luria Broth cultures supplemented with ampicillin [100 □g/mL] inoculated with a single colony (unless otherwise noted) using the GeneJET Plasmid Miniprep Kit as per the manufacturer (Thermo Fisher Scientific, Waltham, Massachusetts).

### Sequencing of plasmid constructs for identity verification and assessment of directionality

The direction and identity of inserts in individual constructs were verified via Sanger sequencing of purified plasmid DNA using M13F, T7, and/or M13-40FOR primers (GENEWIZ from Azenta Life Sciences, Waltham, Massachusetts). To analyze proportions of insert directionality during ligation and within transformant populations, three planarian genes were amplified with their respective primers (SMEST023171001.1: 5’-GTTCACTGGCAGTTTGATTGG-3’ and 5’-CATTTCCTCTTGGCTTGATTGG-3’; SMEST004045004.1: 5’-GCGAACTAACGGGAACAAAC-3’ and 5’-GGCCATGTGCTGGAATAATG-3’; and SMEST012773001.1: 5’-AGGTTAACTGGTGATGCTACTG-3’ and 5’-CCTCCTTTCTAGCTCTGTCTAATTC-3’) and amplicons were processed as specified above. Then, 1 □L of either the ligation reaction or plasmids purified from a mixture of all the white colonies from the transformation were used as template for PCR amplification using primers flanking the multiple cloning site of pGEM-T (5’-GCGCGAATACCTCACTAAGTATACGACTCACTATAGG-3’ and 5’ CGCGCGCTAATACGACTCACTATGATTAGGTGACACTATAG-3’). Amplicons were purified using DNA Clean & Concentrator-5 kit (Zymo Research, Tustin, California), eluted in 15 □L of nuclease-free water and sent for PCR-EZ Sequencing (GENEWIZ, Azenta Life Sciences, Waltham, Massachusetts). FASTQ files from PCR-EZ sequencing were mapped to their respective predicted insertion sequence if inserted sense to the T7 promoter using minimap2 and the command “minimap2 -ax splice -k14 --MD /path/to/reference.fasta path/to/sequenced-results.fastq -o path/to/aligned-sequence.sam” (Li, 2018). The orientation of reads was then quantified using samtools to identify reads sense to T7 promoter with the command “samtools view -c -F 16” and reads sense to the SP6 promoter “samtools view -c -f 16” (Danecek et al., 2021).

### Statistical Analysis

To test for statistical significance in observed distributions of insert orientation for pGEM-T constructs generated in the CUREs, the fraction of constructs with inserts in the sense orientation relative to the T7 promoter were calculated and compared to those generated with sense orientation relative to the SP6 promoter (Figure 1B, 1D). Standard unpaired, two-tailed, Student’s *t*-tests were used to measure statistical significance between the fraction of constructs obtained across the four projects and three genes analyzed. Distribution of orientation for individual projects and cloning events following ligation vs. bacterial selection were tested for statistical significance using chi-squared tests (Figure 1B, 1C, 1D).

### Reagents

| **Reagent** | **Source** | **Cat No/Ref** |
| --- | --- | --- |
| DNA Clean & Concentrator-5 | Zymo Research | D4004 |
| GeneJET Plasmid Miniprep Kit | Thermo Fisher Scientific | K0503 |
| GoTaq 2-Step RT-qPCR System | Promega | A6010 |
| GoTaq® Long PCR Master Mix | Promega | M4021 |
| pGEM®-T Vector Systems | Promega | A3600 |
| TRIzol™ Reagent | Thermo Fisher Scientific | 15596026 |
| *Schmidtea mediterranea*,<br>hermaphroditic strain | Phillip Newmark Laboratory,<br>University of Wisconsin–Madison | (Zayas et al.,<br>2005) |

## Acknowledgements

The authors would like to thank the hundreds of undergraduate students who participated in Molecular Biology laboratory CUREs at UMass Boston and Wright State University over the years. V.D. was supported in part by Graduate Teaching Assistantships and from the University of Massachusetts Boston. VD was additionally supported in part by a College of Science and Mathematics Dean’s Doctoral Research Fellowship with funding from Oracle, project ID R20000000025727. This work was supported by NIH Award No. R15HD082754 to L.R.

